# RNase reverses segment sequence in the anterior of a beetle egg (*Callosobruchus maculatus*, Coleoptera)

**DOI:** 10.1101/059162

**Authors:** Jitse M. van der Meer

## Abstract

The genetic regulation of anterior-posterior segment pattern development has been elucidated in detail for *Drosophila*, but it is not canonical for insects. A surprising diversity of regulatory mechanisms is being uncovered not only between insect Orders, but also within the Order of the Diptera. This raises the question whether the same diversity of regulatory mechanisms exists within other insect Orders. This paper draws attention to the promise of the pea beetle *Callosobruchus maculatus* for elucidating the evolution of pattern regulation mechanisms in Coleoptera and other insect Orders. Introduction of RNase in eggs of *Callosobruchus* replaces anterior segments with posterior segments oriented in mirror image symmetry to the original posterior segments (double abdomens). Reversal is specific for RNase activity, for treatment of the anterior egg pole and for cytoplasmic RNA. Yield depends on developmental stage, enzyme concentration and temperature. A maximum of 30% of treated eggs reversed segment sequence after puncture in 10.0 μg/ml RNase S reconstituted from S-protein and S-peptide at 30 °C. This result sets the stage for an analysis of the genetic regulation of segment pattern formation in the long germ embryo of the Coleopteran *Callosobruchus* and for comparison with the short germ embryo of the Coleopteran *Tribolium*.

## 1. Introduction

In insects, abnormalities in segment sequencing are used to explore the normal regulation of body pattern development along the head-to-tail axis. Partial reversal of segment sequence is among such abnormalities. One type of partial reversal involves the replacement of head and thoracic segments with abdominal segments oriented in mirror image symmetry to the original posterior segments (double abdomen). The converse arrangement occurs in double cephalons. In *Drosophila* the mutations *bicaudal* (Bull 1966) and *bicoid* (Frohnhöfer et al. 1986) produce double abdomen-like arrangements while *dicephalic* produces double heads (Lohs-Schardin 1982). In *Chironomus samoensis* only double abdomen occurs as a mutation (Kalthoff and Albetieha 1986). No double abdomen mutation is known in *Chironomus riparia*, but transcript expression profiling of AP bisected early embryos has revealed a second gene – *panish* – with the same function as *bicoid* in *Drosophila* (Klomp et al. 2015).

Both types of reversal have also been obtained by other experimental means. In the Dipteran *Chironomus* production of double abdomens and double cephalons is correlated with the rearrangement of ooplasm produced by centrifugation of the egg (Yajima 1960). Moreover, in *Chironomus* (Yajima 1964) and *Smittia* (Kalthoff 1971, 1983) UV irradiation of the anterior egg half at the absorption maximum of pyrimidines and purines (253.7 nm) produced double abdomens. UV exposure of the posterior produced double cephalons only in *Chironomus* (Yajima 1964). Development of double abdomens after puncture of the anterior egg of *Smittia* was correlated with the occurrence of an extraovate suggesting the inactivation or elimination of an active component (Schmidt et al. 1975). Puncture of the anterior in RNase produced double abdomens (Kandler-Singer et al. 1976). Together, these results offered the first indication of a role for cytoplasmic polynucleotides in insect segment pattern formation. In sum, experimentally, both types of reversal have been obtained in the Dipterans.

This conclusion is possible because exposure of the anterior or the posterior of these insect eggs to UV light eliminates nuclei as targets. During early development of most insect eggs the dense yolk prevents cell division, but allows division of nuclei which starts with the zygote nucleus. It is located far away from the egg poles in the central yolk mass. Nuclei slowly migrate from this central location in the egg throughout the yolk eventually reaching the yolk-free surface cytoplasm where they are enclosed by cell membranes. Thus early UV treatment localized to the anterior or posterior of the syncytial stage egg cannot affect nuclei because they have not yet arrived there. Nuclei cannot be excluded as targets after puncture in RNase because it can diffuse from the puncture site into the cytoplasm and reach the migrating nuclei.

In Coleoptera only double abdomens have been induced by temporary constriction of eggs of three species: the bean beetles *Callosobruchus maculatus* and *Bruchidius obtectus* and the potato beetle *Leptinotarsa decemlineata* (van der Meer 1984, 1985). The pea beetle *Callosobruchus maculatus* is a global pest of stored legumes. Temporary separation of the yolk mass before nuclei enter the anterior egg half produces a decaying anterior yolk mass (van der Meer 1979: Fig1). This is often accompanied by partial reversal of segment sequence in the posterior egg half provided it is exposed to the decaying anterior fragment. No reversal occurred in the posterior fragment when the anterior egg fragment developed into an anterior half of the embryo (van der Meer 1984). This suggests that reversal in the posterior fragment of the egg is not due to a lack of contact with a developing anterior half, but to a factor leaking from the decaying anterior egg fragment into the posterior one. Since anterior puncture of *Smittia* eggs submerged in RNase caused double abdomens (Kandler-Singer et al. 1976), I test here the hypothesis that in *Callosobruchus* RNase leaking from the decaying anterior egg fragment induced double abdomens in the posterior one.

The purpose of these experiments is to increase the diversity of species for comparative studies of the genetic regulation of anterior-posterior segment pattern development. This regulation has been elucidated in detail for *Drosophila* (Jaeger 2011), but it is not canonical for insects. A surprising diversity of anterior-posterior regulatory mechanisms is being uncovered not only between insect Orders (Lynch 2012; Lynch 2014; Patel 1994; Patel et al. 1994; Sander 1994), but also within the Order of the Diptera (Klomp et al. 2015; Lemke et al. 2008; Lemke and Schmidt-Ott 2009; Lemke et al. 2010; Stauber et al. 2002). This raises the question whether the same diversity of regulatory mechanisms exists within other insect Orders.

The large Order of beetles (Coleoptera) is particularly interesting for comparative embryological study because it includes long, intermediate and short germ types of embryonic development. In long germ development the posterior segment pattern is specified nearly simultaneously with the other segments. Thus, maternal control encompasses the entire segment pattern. By contrast, in short germ development the posterior segments are specified sequentially. Moreover, in many insects with short germ development, the germ rudiment is specified in the posterior region of the egg, far away from the anterior pole. Thus, maternal control covers extra-embryonic, cephalic and sometimes thoracic domains, but excludes abdominal segments which are produced later within the “growth zone”.

It is not known whether the difference in Coleoptera between sequential segment specification in short germ rudiments and simultaneous segment specification in long germ rudiments is correlated with differences in the maternally provided transcription factors involved in these different kinds of specification. Maternally located mRNA transcripts have been identified in the anterior of *Tribolium* eggs (Fu et al. 2012; Schmitt-Engel et al. 2012; Schopfmeier et al. 2009) as well as in the posterior (Schmitt-Engel et al. 2012). It is conceivable that the size of the germ rudiment relative to egg size is correlated with properties of the *maternal* components of segment specification. Therefore, it would be of interest to compare the maternal contributions to the genetic regulation of segment specification within the Coleoptera between the short germ species *Tribolium* and the long germ species *Callosobruchus* (Lynch et al. 2012).

Coleoptera are also suitable for comparative embryological study because genetic regulation of anterior-posterior segment patterning in the short germ embryo of *Tribolium* has been explored to some extent. So far, early maternal and zygotic contributions to establishing the expression domains of zygotic gap, pair-rule and segment polarity genes have been found for the anterior (Bucher et al. 2005; Fu et al. 2012; Kotkamp et al. 2009; Schmitt-Engel et al. 2012; Schoppmeier and Schroeder 2005; Schoppmeier et al. 2009; Schröder 2003; van der Zee et al. 2005; Wolff et al. 1998), the posterior (Copf et al. 2004; Lynch et al. 2012; Schmitt-Engel et al. 2012) as well as for the termini (Schopmeier and Schroder 2005). Pair-rule and segment polarity genes have been identified in two other Coleoptera, *Dermestes* and *Callosobruchus* (Patel et al. 1994). Further study of *Callosobruchus* is promising for a more comprehensive comparison with *Tribolium*.

So far, maternal contributions to segment patterning in *Callosobruchus* have not been explored. Here I provide evidence that RNase activity in the anterior egg can induce double abdomen development suggesting a role for maternal messengers. This sets the stage for using transcript expression profiling (Klomp et al. 2015) of bisected *Callosobruchus* eggs to identify the gene or genes that produce the axis determinants. I also introduce a way of mass collecting eggs.

## 2. Materials and Methods

### 2.1 Adults

Pea beetles (*Callosobruchus maculatus* Fabr., syn. *Bruchus quadrimaculatus* Fabr.) were reared, eggs collected and larval cuticles prepared and analysed according to van der Meer (1979). To avoid development of allergy against pea beetle dust, the incubator in which beetles are reared was placed outside of the working space. Oviposition and egg collection were done at 70 % relative humidity in a hospital baby incubator on a cart. The incubator was fitted with an exhaust fan to create negative internal pressure and an exhaust hose to the outside of the building. Humidity was maintained via an intake from a heated water bath.

### 2.2 Mass collection of eggs on methylcellulose-coated beans

Large numbers of eggs were collected in minutes by letting beetles lay eggs on brown beans coated with methylcellulose. Methylcellulose is a water-soluble carbohydrate polymer produced by Dow Chemical under the brand name Methocel. Grade A indicates methylcellulose products, grade K hydroxypropyl methylcelluloses. Grade A15 or K35 are suitable for making a thin film on beans because their low molecular weight allows for making a low-viscosity solution (Dow Chemical, n.d.).

A 0.5% solution of methocel was prepared by adding 15 °C water to methocel powder very slowly and in that sequence while agitating with a magnetic stirrer until the powder has dissolved. A pouch of flexible nylon mesh containing beans is lowered into a beaker with methocel solution and stirred gently for one hour. The pouch with beans was transferred into another beaker with water. The beans were dried on a screen at room temperature or at 40 °C overnight making sure the beans do not stick to each other. In a dry atmosphere the film is not sticky, but it may become sticky in a humid atmosphere or when touched with fingers.

Eggs were collected according to van der Meer (1979), but on methylcellulose coated beans. Egg-covered beans were placed in a pouch of flexible nylon mesh and the pouch was placed in a beaker with water. The water was kept at 15 °C to slow development and because methylcellulose is more soluble at lower temperature. The pouch was gently moved up and down for 30 minutes. Most eggs detached in 5 minutes and are transferred to 23-24 °C. Under these conditions 98% of detached eggs (n = 44) reached the caudal plate stage and 85% developed into normal larvae (n = 195). The causal plate stage looks normal in the dissecting microscope indicating that a brief exposure to 15 °C does not block segment specification or morphogenesis.

### 2.3 Staging of development

Developmental stages were determined by inspection of serially sectioned eggs (Table 7). Eggs were fixed on the bean in a mixture of 80% ethanol, 40% formaldehyde and undiluted acetic acid in a ratio of 17:1:2 by volume for 20 - 30 minutes at 60 °C and allowed to cool. Eggs were removed from the bean with a razor blade. The chorion was removed from the egg with sharp tungsten needles followed by fixation overnight in the same mixture at room temperature. Eggs can be stored indefinitely in 70% ethanol. Before further processing eggs were washed in 80% ethanol. To spot an egg during and after embedding in paraplast, eggs were stained for 30 to 60 minutes by adding a few drops of thionine to the container with eggs in 80% ethanol (dissolve 1 g. thionine in 10 ml. 100% ethanol at room temperature and add distilled water up to 100 ml.). Eggs were dehydrated via ethanol and xylol and embedded by gradually adding paraplast (Sherwood 60 °C) to the xylol. Eggs were oriented in the liquid paraplast with a hair loop under a dissection microscope. Serial sections – transverse or sagittal – were stained on glycerine coated slides. The glycerine keeps the yolk attached to the slide. Sections were stained with Geidies’ modification of azan-novum (Schulze and Graupner 1960).

### 2.4 Chemicals

RNase A (type III-A from bovine pancreas, EC 3.1.27.5), oxidized RNase A (type XII-AO), RNase S (from bovine pancreas, Grade XII-S), S-peptide (Grade XII-PE), and S-protein (Grade XII-PR) were obtained from Sigma (St. Louis, Missouri), DNase I (from bovine pancreas, deoxyribonucleate 5’-oligonucleotidohydrolase EC 3.1.4.5, Miles Laboratories, Elkhart, Indiana) and Proteinase K (from *Tritirachium album*, EC 3.4.21.64, Boehringer Mannheim, 15686, Indianapolis, Indiana). Oxidized RNase A was obtained by performic acid treatment of RNase A (Richards and Vithayathil 1959). Subtilisin treatment splits RNase A into S-protein and S-peptide that remain attached. This complex is referred to as RNase S and has full RNase activity. S-protein has some residual activity, but S-peptide is inactive (Richards and Vithayathil 1959). All solutions were prepared in distilled water to facilitate diffusion down a concentration gradient into the egg (Kandler-Singer and Kalthoff 1976). DNase I was dissolved immediately before use because solutions containing 100 μg/ml in dilute buffer are stable in the range of pH 4.0 to 9.0 only for a week or longer at about 5°C (Kunitz 1950). RNase S’ (1 μg/ml) was reconstituted by mixing equimolar concentrations of S-peptide (0.84 μg/ml) and S-protein (0.16 μg/ml) (Kandler-Singer and Kalthoff 1976).

### 2.5 Puncturing

Injection into the egg is not possible due to the thickness and tough structure of the chorion. Therefore, eggs were punctured with a sharp glass needle while submersed in various RNases and control substances. The chorion is transparent and single yolk globules are visible. Their displacement shows that the influx of RNase is localized to the puncture site from where it may diffuse. To penetrate the thick chorion it is crucial to use a closed glass needle with sufficient wall thickness. I used type 7740 borosilicate glass with inner filament, outer diameter 1.2 mm, inner diameter 0.6 mm, wall thickness 0.3 mm from Karl Hilgenberg Glaswarenfabrik, Strauchgraben 2, 34323 Malsfeld, Hessen, Germany, www.hilgenberg-gmbh.de. Needles with a diameter tapering to 10 μm were pulled with a Kopf pipette puller.

Eggs were fastened by inserting them with the posterior end into a thin line of soft silicon rubber (Terostat-55 from Henkel Teroson GmbH, Postbox 105 620, Hans-Bunte Strasse 4, 69046 Heidelberg 1, Germany, www.henkel-adhesives.de or www.teroson.de) deposited on a microscope slide. The acetic acid released during hardening of the silicon rubber was rinsed off before puncture.

### 2.6 Classification and calculation of results

‘Dead’ embryos includes all embryos that did not produce a larva with a differentiated cuticle. ‘Normal’ embryos comprises embryos that produced larvae with a normal segment pattern including one or two with missing anterior segments due to puncturing damage. Percentages are calculated as a fraction of punctured eggs.

## 3 Results

### 3.1 Controls

Nuclear migration stage eggs were punctured at the anterior pole while submerged in distilled water (pH 6) and solutions of DNase I, oxidized RNase A, proteinase K, S-protein and S-peptide, all at 23-24 °C (Table 2). No control treatment produced reversal except S-protein (8% reversal at 5 μg/ml and 9% at 10.0 μg/ml) in accordance with its residual RNase activity (Richards & Vithayathil, 1959: Fig. 3). RNase S’ which is reconstituted from S-protein and S-peptide in equimolar amounts produced 11% reversal at 5.0 μg/ml and 24% at 10.0 μg/ml. (Table 4, Fig1a) confirming that the combination of S-protein and S-peptide has higher RNase activity than S-protein alone. These results indicate that partial reversal of segment sequence is not due to the effect of solvent (distilled water), enzyme activity not specific for RNA (DNase I, proteinase K) or of protein (oxidized RNase A, S-peptide). I conclude that partial reversal of segment sequence is specific for RNase activity.

**Table 1:** Dependence of developmental stages on temperature (°C). Development times are in hours ± 15 minutes. Development slows down when eggs develop in water or oil. SBy: nuclei have just arrived in periplasm, no transverse cell membranes, pole cells formed; SBm1: transverse cell membranes begin to form; Sbm2 & SBo: transverse cell membranes completed; CBy: tangential cell membranes completed; CBo: early ventral plate.

| Stage description | Abbreviation | 70% relative humidity |  |  | distilled water |  |  |
| --- | --- | --- | --- | --- | --- | --- | --- |
|  |  | 30 | 25 | 22 | 30 | 25 | 22 |
| meiosis I - anaphase | M I | 0.25 | 0.5 | 1-3.5 | — | — | 4 |
| meiosis II - fusion pronuclei | M II | 1 | 1.5 - 2.5 | 4 | — | — | — |
| 2 nuclei | NM 2 | 2 | 3 - 4 | 5 - 6 | — | 4 | 8 |
| 4 nuclei | NM 4 | — | 4 | 6 - 7 | — | — | — |
| 4-8 nuclei | NM 4-8 | 3 | — | — | — | — | — |
| 8-16 nuclei | NM 8-16 | — | 5 - 6 | 8 | — | — | — |
| 16-32 nuclei | NM 16-32 | 4 | — | 8 | 4 | — | — |
| 32-64 nuclei | NM 32-64 | 5 | 6 - 7 | 9 - 10 | — | — | — |
| 64-128 nuclei | NM 64-128 | — | 8 | 10 - 11 | — | — | 12 |
| 128-256 nuclei | NM 128-256 | 6 | — | 11 - 12 | — | 8 | — |
| young syncytial blastoderm | SBy | 6 | 9.5-10.5 | 12 - 13 | 8 | 12 | — |
| mid-syncytial blastoderm 1 | SBm1 | 8 | — | 16 | — | — | 24 |
| mid-syncytial blastoderm 2 | SBm2 | 10 | 12 | 20 | — | 16 | — |
| old syncytial blastoderm | SBo | 12 | — | 24 | — | — | — |
| young cellular blastoderm | CBy | 13 | 16 | — | — | — | — |
| old cellular blastoderm | CBo | 14 | 18 - 20 | — | — | 20 | — |

**Table 2:**
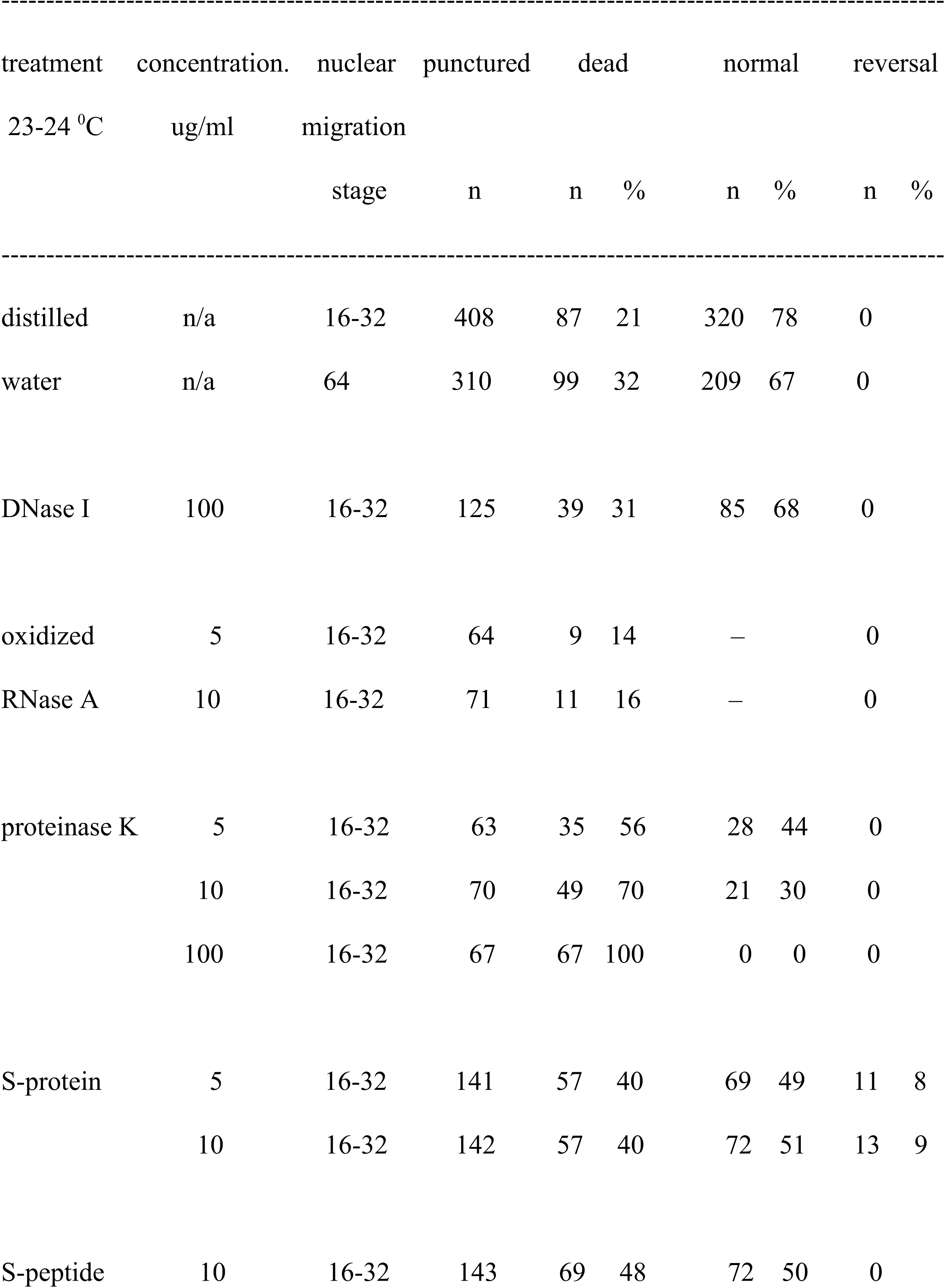
Controls

**Figure 1.**
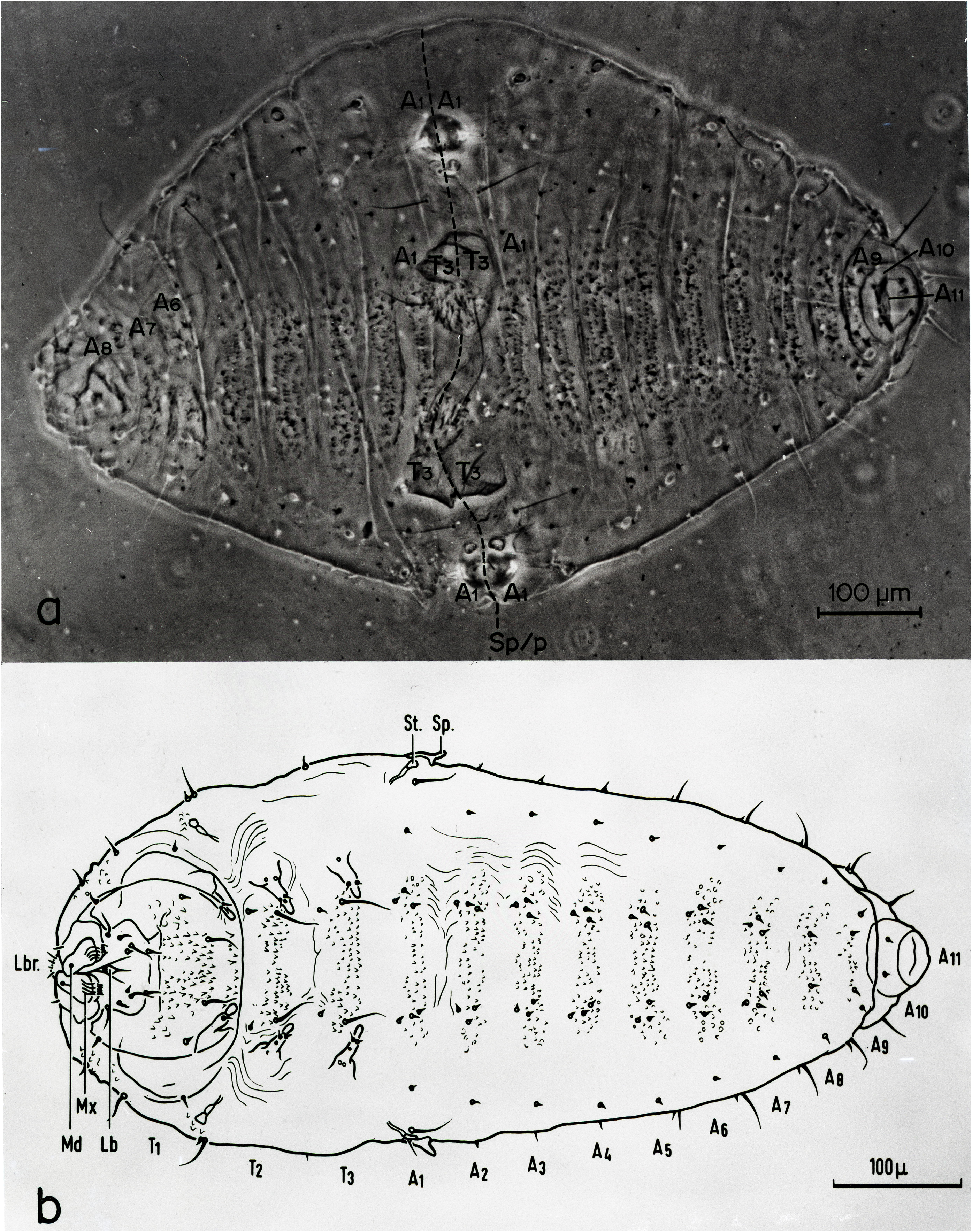
Double abdomen and normal first instar larvae of *Callosobruchus maculatus*. Ventral views of (a) double abdomen with reversed second abdomen pointing left (phase contrast), (b) normal larva. Comparison shows that in (a) the head segments and the first two thoracic segments have been replaced by a reversed sequence of T3 through A8 which joins the non-reversed abdomen located to the right of the dashed symmetry line (Sp/p). Abbrev.: head segments: Lbr: labrum, antennae (not visible), Md: mandibulae, Mx: maxillae, Lb: labium; thorax: 3 segments T1, T2, and T3, each with a pair of legs (L); abdomen: A1– A11, St: stigma, Sp: spine of segment A1 (reproduced with permission by Taylor & Francis Ltd. www.taylorandfrancis.com from van der Meer 1985 Fig1)

### 3.2 Stage dependence of reversal of segment sequence

To determine whether the frequency of partial reversal of segment sequence depends on stage of development, eggs were punctured at the anterior pole while submerged in 0.5 μg/ml RNase A. There is a well-defined dependence on stage of development with a maximum reversal yield of 14% of punctured eggs at nuclear migration stages between 16 and 64 nuclei (Table 3).

**Table 3:** Influence of stage of development on the frequency of reversal after anterior puncture in 0.5 μg/ml RNase A.

| stage |  | punctured |  | dead |  | normal |  | reversal |  |
| --- | --- | --- | --- | --- | --- | --- | --- | --- | --- |
| (hr at 30 °C) |  | n |  | n | % | n | % | n | % |
| NM2 | (2hr) | 171 |  | 61 | 36 | 103 | 60 | 7 | 4 |
| NM4-8 | (3hr) | 236 |  | 94 | 40 | 125 | 52 | 17 | 7 |
| NM16-32 | (4hr) | 184 |  | 46 | 25 | 112 | 61 | 26 | 14 |
| NM64 | (5hr) | 183 |  | 59 | 32 | 99 | 54 | 25 | 14 |
| SBy | (6hr) | 166 |  | 42 | 25 | 115 | 69 | 9 | 5 |
| SBm | (7hr) | 188 |  | 51 | 27 | 137 | 73 | 0 | 0 |

### 3.3 Concentration dependence of reversal of segment sequence

In order to increase the yield of double abdomens I determined the effect of enzyme concentration after anterior puncture at the stage that yielded a maximum frequency of partial reversal of segment sequence (NM16-32, Table 3). While the death rate varies considerably between concentrations, it tends to decline at higher enzyme concentration (Table 4). There is a well-defined concentration-dependence with a maximum yield of 22% double abdomens at 2.0 μg/ml RNase A and 17% at 1.0 μg/ml of RNase S (Table 4).

**Table 4:** Influence of enzyme concentration on reversal yield after anterior puncture in RNase A and RNase S at NM16-32

| RNase A |  |  |  |  |  |  |  | RNase S |  |  |  |  |  |  |
| --- | --- | --- | --- | --- | --- | --- | --- | --- | --- | --- | --- | --- | --- | --- |
| µg/ml | punctured | dead |  | normal |  | reversal |  | punctured | dead |  | normal |  | reversal |  |
|  | n | n | % | n | % | n | % | n | n | % | n | % | n | % |
| 0.2 | 152 | 20 | 13 | 132 | 87 | 0 | 0 | — | — | — | — | — | — | — |
| 0.5 | 387 | 112 | 29 | 229 | 59 | 46 | 12 | 139 | 83 | 60 | 56 | 40 | 0 | 0 |
| 0.8 | 132 | 77 | 58 | 37 | 28 | 18 | 14 | 143 | 83 | 58 | 58 | 41 | 2 | 1 |
| 1.0 | 283 | 134 | 47 | 114 | 40 | 35 | 12 | 312 | 68 | 22 | 190 | 61 | 54 | 17 |
| 2.0 | 143 | 55 | 38 | 57 | 40 | 31 | 22 | 143 | 71 | 50 | 67 | 47 | 5 | 3 |
| 5.0 | 138 | 120 | 87 | 9 | 7 | 9 | 7 | 136 | 88 | 65 | 29 | 21 | 19 | 14 |
| 8.0 | 134 | 128 | 96 | 4 | 3 | 2 | 2 | 140 | 108 | 77 | 15 | 11 | 17 | 12 |

RNase S’ is reconstituted from S-protein and S-peptide. It is included as a standard for comparison against the control test with S-protein and S-peptide (Table 2). I also determined dependence of yield on the concentration of RNase S’ to see whether a higher yield could be obtained (Table 5). RNase S’ produced 24% double abdomens at 10.0 μg/ml compared to RNase A which produced 22% at 2.0 μg/ml.

**Table 5:** Reversal yield after anterior puncture in RNase S’ at NM16-32

| $\mu\text{g/ml}$ | punctured | dead | | normal | | reversal | |
| --- | --- | --- | --- | --- | --- | --- | --- |
|  | n | n | % | n | % | n | % |
| 1.0 | 136 | 24 | 18 | 110 | 81 | 2 | 2 |
| 2.0 | 131 | 87 | 66 | 29 | 22 | 15 | 12 |
| 5.0 | 195 | 127 | 65 | 46 | 24 | 22 | 11 |
| 10.0 | 131 | 42 | 32 | 57 | 44 | 32 | 24 |

### 3.4 Influence of puncture site on reversal of segment sequence

Syncytial stage eggs were punctured during early nuclear migration NM 16-32 while submerged in 0.5 μg/ml of RNase A (23-24 °C). The puncture site is precisely controlled using the shape of the egg which looks like a chicken egg cut in half lengthwise. Thus, the anterior of the egg is round and is identified by the location of the head of the embryo later in development. The posterior is pointed and is identified by the location of the abdomen later on. The ventral side is dome-shaped and is identified by where later the embryo develops. The dorsal side is flat and is the location where extra-embryonic tissue develops later on (van der Meer 1979, Figs. 1, 7). On average eggs are 700 μm long, 200 μm wide and 150 μm largest height. Lateral puncture was on the left at 50% of egg length, midway between the flat dorsal side and the domed ventral side of the egg. Ventral and dorsal puncture was at 50% of egg length half way between the sides of the egg. These puncture sites are close to the zygote nucleus which is located at 60 ± 10% of egg length equidistant from the surface of the egg.

The survival rate after mid-dorsal puncture was extremely low. This cannot be due to the loss of the maternal pronucleus in the maturation island which is on the dorsal side because these eggs were punctured during early nuclear migration.

### 3.5 Effect of temperature on frequency of reversal of segment sequence

The correlation between reduced RNase activity in protein S and lower reversal frequency (Table 2) supports the conclusion that reversal is the specific result of the level of RNase activity. To further support that correlation, and also to maximize reversal yield for future experiments, nuclear migration stage eggs with 64 nuclei were submitted to 30 °C before, during and after anterior puncture in 10.0 μg/ml RNase S’ (Table 7). At this concentration NM 16-32 eggs punctured at room temperature (23-24 °C) produced 24% reversal (Table 5). Following oviposition eggs were kept for 5 hrs at 30 °C, that is up to NM 64. Puncturing lasted from 5 to 6 hrs at 30 °C followed by incubation at 30 °C. Control eggs underwent the same temperature regimens as the punctured eggs, but were not attached to adhesive or punctured.

At 30 °C, RNase redirected slightly less than half of the eggs that developed normally after puncture in distilled water (68%) to partially reversing their segment sequence (30%) without affecting the survival rate which is 67% and 68%, respectively (Table 7: top). From there the reversal frequency declined with temperature to 24% at 23-24 °C (Table 7).

In sum, reversal yield depends on developmental stage, enzyme concentration and temperature. A maximum of 30% of eggs treated at 30 °C with 10.0 μg/ml RNase S reconstituted from S-protein and S-peptide developed double abdomens.

## 4. Discussion

Introduction of RNase into the anterior of the egg of *Callosobruchus* replaces anterior segments with posterior ones in mirror image symmetry to the original posterior segment pattern (double abdomen). Reversal is specific for RNase activity because inactivated RNase, proteinase and DNase did not produce double abdomens. Thus reversal is not due to an effect on protein, but on RNA. RNase treatment does not identify the species of RNA involved in reversal.

The strongest evidence for the specificity of the effect is that at the same concentration (10 μg/ml) and temperature (23-24 °C) the frequency of reversal was correlated with the degree of RNase S’ activity. S-peptide has no residual RNase activity and produced no reversal. S-protein has residual RNase activity and produced 9% reversal. RNase S’ which is reconstituted from S-protein and S-peptide produced 24% reversal. These observations indicate that the partial reversal of segment sequence is due specifically to the enzymatic destruction of RNA.

The rise of reversal frequency from 24% at 23-24 °C to 30% at 30 °C (Table 7) supports the conclusion that reversal is due to RNase activity since it depends on temperature. But this rise could also be due in whole or in part to an effect of temperature directly on survival frequency. Eggs may be more sensitive to puncture at lower temperature due to the delayed healing of injection damage. One would expect a lower survival rate after puncture in distilled water at 23-24 °C than at 30 °C. But the survival rate was higher at lower temperature (78%) than at higher temperature (68%) (Table 7). Therefore, the rise in reversal frequency can be attributed to the effect of temperature on RNase activity rather than on survival rate.

The regular rise of reversal frequency with RNase concentration also suggests that reversal is due to RNase activity. Reversal frequency tends to rise regularly with enzyme concentration up to a maximum after which it declines (Tables 4, 5). This can be explained by assuming that above an optimum concentration the enzyme kills the embryo. Thus the death rate should rise with enzyme concentration and it tends to do so. But it varies widely though not irregularly between enzyme concentrations for both RNase A and RNase S (Tables 4, 5) as well as for RNase S’ (Table 6). For instance, the death rate declines from 58% in 0.8 μg/ml RNase A to 38% in 2.0 μg/ml, then jumps to 96% in 8.0 μg/ml RNase A (Table 4). Further, when the death rate rises the frequency of normal embryos should decline. Again, this is the overall tendency, but individual frequencies vary widely between enzyme concentrations. For instance, the frequency of normal development declines regularly with increasing concentrations of RNase A over seven concentrations, but jumps from 28% to 40% between 0.8 μg/ml and 1.0 μg/ml (Table 4). This variation is not due to differences in quality between egg batches because eggs are not deposited in batches, but individually. The effect of a particular concentration was determined in two, sometimes three, experiments run on consecutive days using the same stock solution. The same variation was observed between experiment runs. I speculate that variation in dilution of enzyme stock solution may explain the variation of frequencies between experiment runs testing the same concentration as well as between enzyme concentrations.

**Table 6:** Influence of puncture site on reversal of segment sequence in 0.5 μg/ml RNase A at NM 16-32.

| site | punctured | dead |  | normal |  | reversal |  |
| --- | --- | --- | --- | --- | --- | --- | --- |
|  | n | n | % | n | % | n | % |
| posterior | 156 | 19 | 12 | 137 | 88 | 0 | 0 |
| lateral | 127 | 33 | 26 | 94 | 74 | 0 | 0 |
| ventral | 119 | 74 | 62 | 45 | 38 | 0 | 0 |
| dorsal | 120 | 119 | 99 | 1 | 1 | 0 | 0 |
| anterior | 387 | 112 | 29 | 229 | 59 | 46 | 12 |

**Table 7:** Influence of temperature before, during and after puncture in 10 μg/ml RNase S’ during early nuclear migration.

| temperature | stage | punctured | dead |  | normal |  | reversal |  |
| --- | --- | --- | --- | --- | --- | --- | --- | --- |
|  |  | n | n | % | n | % | n | % |
| 30 °C | NM64 | 138 | 46 | 33 | 50 | 36 | 42 | 30 |
| control (puncture<br>in distilled water) | NM64 | 310 | 99 | 32 | 211 | 68 | 0 | 0 |
| 23-24 °C | NM16-32 | 131 | 42 | 32 | 57 | 44 | 32 | 24 |
| control (puncture<br>in distilled water) | NM16-32 | 408 | 87 | 21 | 320 | 78 | 0 | 0 |

In *Drosophila* (Diptera), *Tribolium* (Coleoptera) and *Nasonia* (Hymenoptera) the sources of the protein gradients that specify the A-P segment pattern (maternal mRNAs) are located in the cytoplasm in the two ends of the egg (Lynch et al. 2012). In *Callosobruchus*, however, both nuclear versus cytoplasmic location and geographic location remain to be clarified. If RNase diffuses from the puncture site into the egg it may reach both nuclear and cytoplasmic targets located away from the puncture site. But a hypothetical involvement of nuclear targets can be excluded. During early nuclear migration NM 16-32 lateral puncture failed to produce reversal (Table 6). Lateral puncture is positioned at 50% of egg length and nuclei spread from the location of the zygote nucleus at 60 ± 10% of egg length. Thus during early nuclear migration NM 16-32 lateral puncture is much closer to the migrating nuclei than anterior puncture which is at 100% of egg length (700 μm). If diffusion of RNase from the anterior puncture site would have produced reversal by affecting remote nuclear targets involved in segment specification so much the more would have diffusion from a lateral puncture site which is closer to the nuclei. But this did not happen (Table 6). Therefore, nuclear targets can be excluded at this stage and this applies also to anterior puncture.

As for the cytoplasmic location of targets, the failure of lateral and posterior puncture to produce reversal (Table 6) also excludes these areas from involvement in segment specification at NM 16-32. Thus, the effect of anterior puncture is specific for the anterior egg region during early nuclear migration NM 16-32. At that stage the targets for RNase associated with A-P segment specification are located in the anterior egg region. No data are available to extend this conclusion to other stages, but it is unlikely that target RNAs involved in A-P segment pattern regulation would be located laterally, ventrally or dorsally.

In conclusion, the data presented do not identify the species of RNA involved in segment specification, but they do justify the conclusion that the target is RNA and that this RNA is not located in the nuclei, but in the anterior egg cytoplasm during early nuclear migration (NM 16-32). These data are consistent with a role for maternally deposited messenger RNA in this process, To mention one hint in this direction, the pair-rule gene *even-skipped* has the same function in *Callosobruchus* (Patel et al. 1994) as in *Drosophila*, namely to mark the anterior margins of odd-numbered *engrailed* stripes. Since *even-skipped* in *Drosophila* is regulated by morphogens produced by maternal messengers located in the egg poles, it may be regulated by maternal messengers in a similar way in *Callosobruchus*.

The purpose of this paper is to draw attention to *Callosobruchus* for elucidating the evolution of pattern regulation mechanisms in insects. Mechanisms for the specification of A-P segmentation have turned out to be very diverse between and within insect Orders (Section 1). The question is whether there are any patterns within this diversity that can clarify how it evolved at the molecular level. The RNA-dependent control of AP segmentation in *Callosobruchus* and the yield of one reversal in every three punctured eggs makes this species a promising candidate for expanding the repertoire of molecular mechanisms of genetic control of segmentation and for elucidating evolutionary relationships. Moreover, the maternal effect genes *causal* and *hunchback* as well as various gap and pair-rule genes have been identified in *Callosobruchus* (Benton et al. 2016). These developments set the stage for identifying the gene or genes regulating segment specification by transcript expression profiling (Klomp et al. 2015) of bisected *Callosobruchus* eggs. This will make possible a comparison of genetic regulation mechanisms within Coleoptera and clarify their evolution.

## Acknowledgments

Research was carried out in the laboratories of Dr. J. M. Denucé, Department of Zoology, Radboud University, Nijmegen, The Netherlands (1978), Dr. W. Müller, Department of Zoology, University of Heidelberg, Germany (1978-79) and Dr. L. F. Jaffe, Department of Biological Sciences, Purdue University, Indiana, USA (1980-82). The results are published only now because of renewed interest in comparing mechanisms of pattern development between insect Orders. This work was supported by a grant from the Commission of the European Union for Research in Molecular Biology and Radiobiology 1978-1979 to Dr. J. M. van der Meer and by National Science Foundation grants to Dr. L. F. Jaffe. I thank Dr. U. Schmidt-Ott for many helpful comments on a previous draft of this paper.

The author declares that he has no conflict of interest.

